# Software-assisted manual review of clinical NGS data: an alternative to routine Sanger sequencing confirmation with equivalent results in >15,000 hereditary cancer screens

**DOI:** 10.1101/305011

**Authors:** Dale Muzzey, Shera Kash, Jillian I. Johnson, Laura M. Melroy, Piotr Kaleta, Kelly A. Pierce, Kaylene Ready, Hyunseok P. Kang, Kevin R. Haas

## Abstract

Clinical genomic tests increasingly utilize a next generation sequencing (NGS) platform due in part to the high fidelity of variant calls, yet rare errors are still possible. In hereditary cancer screening, failure to correct such errors could have serious consequences for patients, who may follow an unwarranted screening or surgical-management path. It has been suggested that routine orthogonal confirmation via Sanger sequencing is required to verify NGS results, especially low-confidence positives with depressed allele fraction (<30% of alternate allele). We evaluated whether an alternative method of confirmation—software-assisted manual call review—performed comparably to Sanger confirmation in >15,000 samples. Licensed reviewers manually inspected both raw and processed data at the batch-, sample-, and variant-level, including raw NGS read pileups. Of ambiguous variant calls with <30% allele fraction (1,707 total calls at 38 unique sites), manual call review classified >99% (1,701) as true positives (enriched for long insertions or deletions (“indels”) and homopolymers) or true negatives (often conspicuous NGS artifacts), with the remaining <1% (6) being mosaic. Critically, results from software-assisted manual review and retrospective Sanger sequencing were concordant for samples selected from all ambiguous sites. We conclude that the confirmation required for high confidence in NGS-based germline testing can manifest in different ways: a trained NGS expert operating platform-tailored review software achieves quality comparable to routine Sanger confirmation.

## INTRODUCTION

Hereditary cancer screening (HCS) has well established clinical validity and clinical utility: it ascribes increased cancer risk to particular germline variants^1–3^, and the presence of these variants often alters patient management^4,5^. As a result, accurate variant detection is critical. Though multiple HCS validation studies demonstrate exemplary analytical sensitivity and specificity^6,7^, even rare analytical errors—false positives or false negatives—could grossly misrepresent an individual patient’s risk of cancer and have grave clinical consequences.

Though NGS is now mature and widely used for HCS^8,9^, in its nascency NGS was particularly susceptible to false positives, generally resulting from low coverage and high rates of both random errors and systematic errors on early instruments^10–13^. For instance, NGS data alone could not resolve the genotype at a site with 4x depth and 25% allele fraction (i.e., one read with an alternate base and three reads with reference bases): either a sequencer error occurred in a patient homozygous for the reference allele, or the allele fraction was depressed in a heterozygous patient due to the discreteness of few observations.

Variant-calling ambiguities in early NGS data prompted medical societies to suggest that variants identified on clinical panels required confirmation^14,15^. With low-depth, high-error NGS data, no level of algorithmic sophistication or added manual scrutiny could resolve certain genotypes; therefore, confirmatory evidence needed to come from an orthogonal technology, typically Sanger sequencing^16^. However, advances in NGS technology over time (e.g., lower per-base error rates and hybrid-capture protocols that yield high depth in regions of interest) have substantially increased achievable variant-calling confidence^12^, calling into question the recommendation for routine confirmation of NGS results. Indeed, whether Sanger sequencing confirmation of NGS data should be routine is a contentious topic in the clinical genomics field: some studies argue that it is critical^17,18^; others suggest that it is largely unnecessary ^19,20^, and yet another claims that it can actually increase the odds of a clinical laboratory returning false results^21^.

In a concordance analysis of NGS and Sanger sequencing data from >20,000 HCS patients, Mu and colleagues suggested that the utility of Sanger sequencing depends largely on the allele fraction of ambiguous variant calls^17^. For calls with allele fraction >30%, orthogonal confirmation was not required to achieve high sensitivity and specificity. However, the authors recommended routine use of Sanger confirmation for putative positive calls with low allele fraction (e.g., <30%) because Sanger sequencing confirmed only a subset of low-confidence NGS positive calls. The authors noted that many variants adjudicated by Sanger sequencing could have been resolved via manual inspection of the NGS data but that such inspection was infeasible and error-prone in a high-throughput laboratory.

At the inception of our clinical laboratory, we engineered a scalable results-management database and software interface wherein expert human reviewers could confirm the quality of assay results prior to reporting. The cloud-deployed and/or LIMS-integrated software—termed “<u>Ma</u>nual <u>C</u>all <u>R</u>eview <u>O</u>ptimization” (MaCRO) and described further herein—now supports multiple different clinical products and assay types (e.g., NGS for HCS, PCR for fragile X carrier status, cfDNA sequencing for noninvasive prenatal screening, etc.), allows viewing of raw and processed data, permits rapid identification of samples requiring retest, enables and records manual variant-call overrides (with comments and auditability) if the expert disagrees with the bioinformatics algorithms, and tracks sample-specific actions and discussions. Critically, the system implements a workflow by which every variant call that could potentially be reportable (e.g., deleterious variants and variants of uncertain significance (VUS) in the context of HCS) is manually confirmed by an expert via the MaCRO interface. Only if the combination of algorithmic assay results and MaCRO cannot confirm a variant call is confirmation pursued through alternate methods such as Sanger sequencing.

Here we evaluated the efficacy of MaCRO confirmation as an alternative to Sanger confirmation to yield confident variant calls in more than 15,000 HCS patients. We find that MaCRO facilitates inspection of raw NGS data underlying each variant call and can confidently disambiguate low-confidence genotype calls (e.g., those with low allele fraction). MaCRO confirmation is fast, with a single operator able to confirm all reportable calls for a batch of nearly 100 samples in 15 minutes on average. Further, MaCRO confirmation is accurate: retrospective Sanger sequencing showed perfect concordance between MaCRO and Sanger results. The accuracy and efficiency of MaCRO confirmation demonstrates that Sanger confirmation is not the only methodology by which laboratories can achieve confident variant calls in HCS.

## METHODS

### Institutional Review Board approval

The study protocol was reviewed and designated as exempt by Western Institutional Review Board (WIRB). Patient information was de-identified according to the Health Insurance Portability and Accountability Act Privacy Rule. An informed consent waiver was approved by WIRB.

### Patient cohort

The study includes variant-calling results from 15,080 de-identified patients who underwent HCS between May 1, 2015 and October 31, 2016. Patients from New York State or from outside the U.S., as well as those who elected to opt out of research, were excluded from the study.

### Screen description

The HCS workflow begins with assembly of a sequencing batch (Figure 1A). DNA from patients’ blood or saliva samples is extracted and prepared for NGS via barcoded adapter ligation and PCR. A 96-well batch contains clinical samples, cell-line samples that act as genotype controls (e.g., NA12878), and no-template wells to detect contamination.

**Figure 1:**
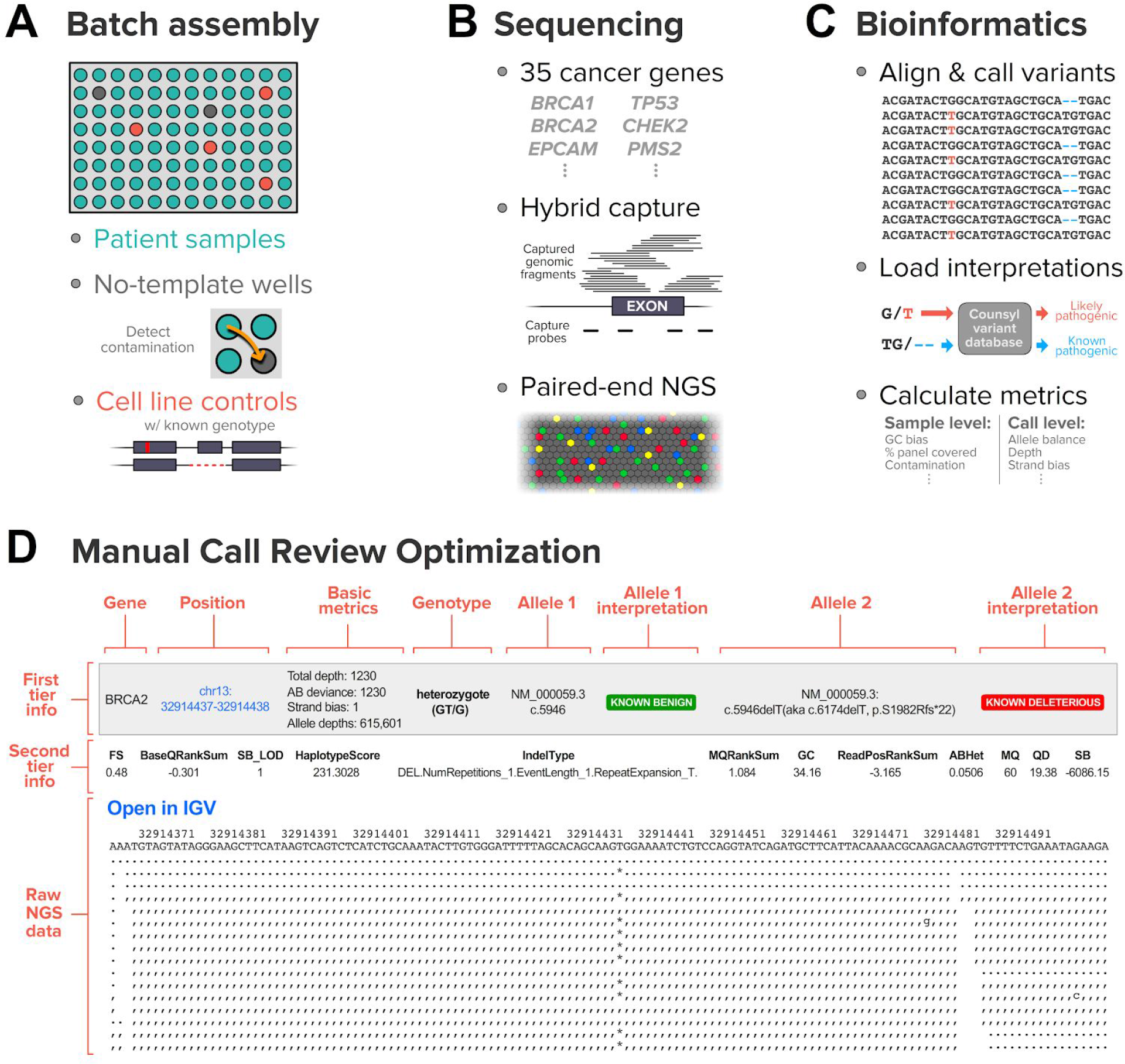
Overview of assay design, quality-control measures, and variant-calling pipeline that populate the manual call review optimization interface. **(A)** Each sequencing batch combines patient samples, cell-line controls, and no-template wells, all in randomized positions. **(B)** Hybrid capture enables targeted sequencing of genes of interest via multiplexed paired-end NGS. **(C)** In the bioinformatics pipeline, alignment precedes variant calling, which is followed by both the loading of interpretations and calculation of calling metrics. **(D)** The MaCRO interface for individual calls renders information in tiers, with deeper tiers revealed to the user upon clicking. Both basic and complex metrics are accessible via the interface, and the raw NGS data centered on the variant of interest is viewable directly in text or via a link to a genome browser.

During the sequencing step (Figure 1B), hybridization-capture probes enrich for targeted regions of interest (20nt padded exons and known-deleterious, deep-intronic sites). In this study, we considered variants in the following genes: *APC*, *ATM*, *BARD1*, *BMPR1A*, *BRCA1*, *BRCA2*, *BRIP1*, *CDH1*, *CDK4*, *CDKN2A*, *CHEK2*, *EPCAM*, *MEN1*, *MLH1*, *MRE11A*, *MSH2*, *MSH6*, *MUTYH*, *NBN*, *PALB2*, *PMS2*, *POLD1*, *POLE*, *PTEN*, *RAD50*, *RAD51C*, *RAD51D*, *RET*, *SMAD4*, *STK11*, *TP53*, and *VHL*. We perform paired-end NGS, described previously elsewhere^6^.

After sequencing, variant calling commences in the bioinformatics pipeline (Figure 1C). SNVs and indels are identified with the Genome Analysis Toolkit (version 1.6 in the study time period)^22^, Freebayes ^23^, and custom genotyping software for representing complex haplotypes spanning clustered calls. Though their quality assessment is not addressed here, copy-number variants (CNVs) are found via a custom calling algorithm^6^ and undergo MaCRO confirmation as well. Prior to rendering in MaCRO, the bioinformatics pipeline computes various metrics at the sample level (e.g., GC bias, fraction covered) and call level (e.g., depth, strand bias, read-position bias, and allele fraction).

### Manual Call Review Optimization (MaCRO)

The MaCRO software interface is a custom database-backed web application implemented in the Django framework leveraging Postgres optimizations. It loads calls and their associated metrics from the bioinformatics pipeline, as well as pathogenicity interpretations (from tools like SNPEFF^24^, from external resources like ClinVar^25^ and dbSNP^26^, and from our internal database). Because it is software driven, the review workflow is strictly controlled and, therefore, robust, auditable, and reproducible from batch to batch and from operator to operator over time.

The first step in the workflow includes evaluation of batch-level metrics (e.g., number of samples passing QC criteria, average sample depth) and confirmation that both control-sample genotypes and QC metrics matched expectation. If metrics or control calls are unexpected, the reviewer can fail the batch to queue it for retesting. Next, the operator reviews sample-level metrics, which include pre-sequencing (e.g., DNA concentration) and post-sequencing (e.g., base quality, depth variance, contamination, etc.) quality-control data. Sample with metrics outside of validated boundaries are queued for retesting.

Upon passing of the batch, controls, and qualified samples, a single secure webpage loads and segments all calls from the entire batch into separate tables for SNVs/indels (Figure 1D) and CNVs. To balance the needs for expedient and meticulous review, the interface displays each call’s information via tiered panels of increasing detail, the last of which depicts raw NGS reads (Figure 1D; the page also links to a genome browser for graphical representation of reads). MaCRO confirmation of variant calls in a batch is performed by a single operator, but any call overrides (i.e., where manual inspection reverses the algorithmically determined genotype) can be flagged and annotated directly within the MaCRO software, prompting review by a second operator to minimize the possibility of human error. Tabs stratify calls based on the variants’ confidence (i.e., call vs. no-call) and clinical-interpretation class (i.e., known deleterious, likely deleterious, VUS, likely benign, and known benign). Every variant in a patient’s ordered panel that is deleterious or a VUS is reviewed. Known- and likely-benign variant calls do not undergo routine review as part of the MaCRO SOP, but they are loaded into the interface for ready access. Finally, the MaCRO software compiles data from multiple runs of a single sample (when applicable) to ensure within-sample concordance prior to reporting.

A MaCRO operator in our laboratory must be licensed as a Clinical Genetic Molecular Biologist Scientist in California and have a minimum of 2 years of experience performing NGS and/or reviewing NGS results. Four operators, who ranged in experience with MaCRO but all underwent training with the software, executed MaCRO on the patient cohort profiled in this study.

For this study, variant calls and their metrics, raw data links (e.g., BAM files), and timestamps were queried from our production databases, de-identified, and then analyzed. Calculations of the time required to perform MaCRO on a batch measured the duration between timestamps of the batch’s first passed control and third-to-last passed test sample (this latter timestamp accounts for samples requiring attention outside of MaCRO; described in Results). Because this proxy for active review time does not capture MaCRO-independent interruptions to the operators’ workflow (e.g., meetings, meals, working hours), batches were not considered if duration exceeded 100 minutes, an upper threshold selected because it conspicuously captures the empirical distribution of MaCRO review time (see Results).

### Sanger confirmation

For Sanger sequencing confirmation of a putative positive call at a given site, the region flanking the site was PCR amplified using DNA extracted from a clinical sample of interest. This amplified genomic DNA was the substrate for bidirectional Sanger sequencing reactions (BigDye, ThermoFisher), which used custom and manually designed sequencing primers that were ~100nt upstream or downstream of the site. Sanger sequencing traces were acquired on a 3730 instrument (ThermoFisher) and interpreted in 4Peaks.

## RESULTS

Our assessment of MaCRO confirmation for all potentially reportable positive calls began with a compilation of each call’s allele fraction and read depth, as these two factors are key drivers of call confidence (Figure 2A). Reportable calls for HCS include known deleterious and likely deleterious variants, as well as VUSs. As expected for a germline test, most positive calls are from heterozygous sites and, therefore, have an allele fraction near 50%: the normally distributed population of clear positives—9,424 calls spread across 3,632 unique sites—had median allele fraction of 49.3% with standard deviation of 4.9%. However, 15.3% of calls (1,707 calls spanning just 38 unique sites) have allele fraction <30%. We term this region the “ambiguous zone” because it is more than four standard deviations from the mean of clear positives in our data, and because it was shown previously to be enriched for NGS variant calls overturned via Sanger sequencing^17^.

**Figure 2:**
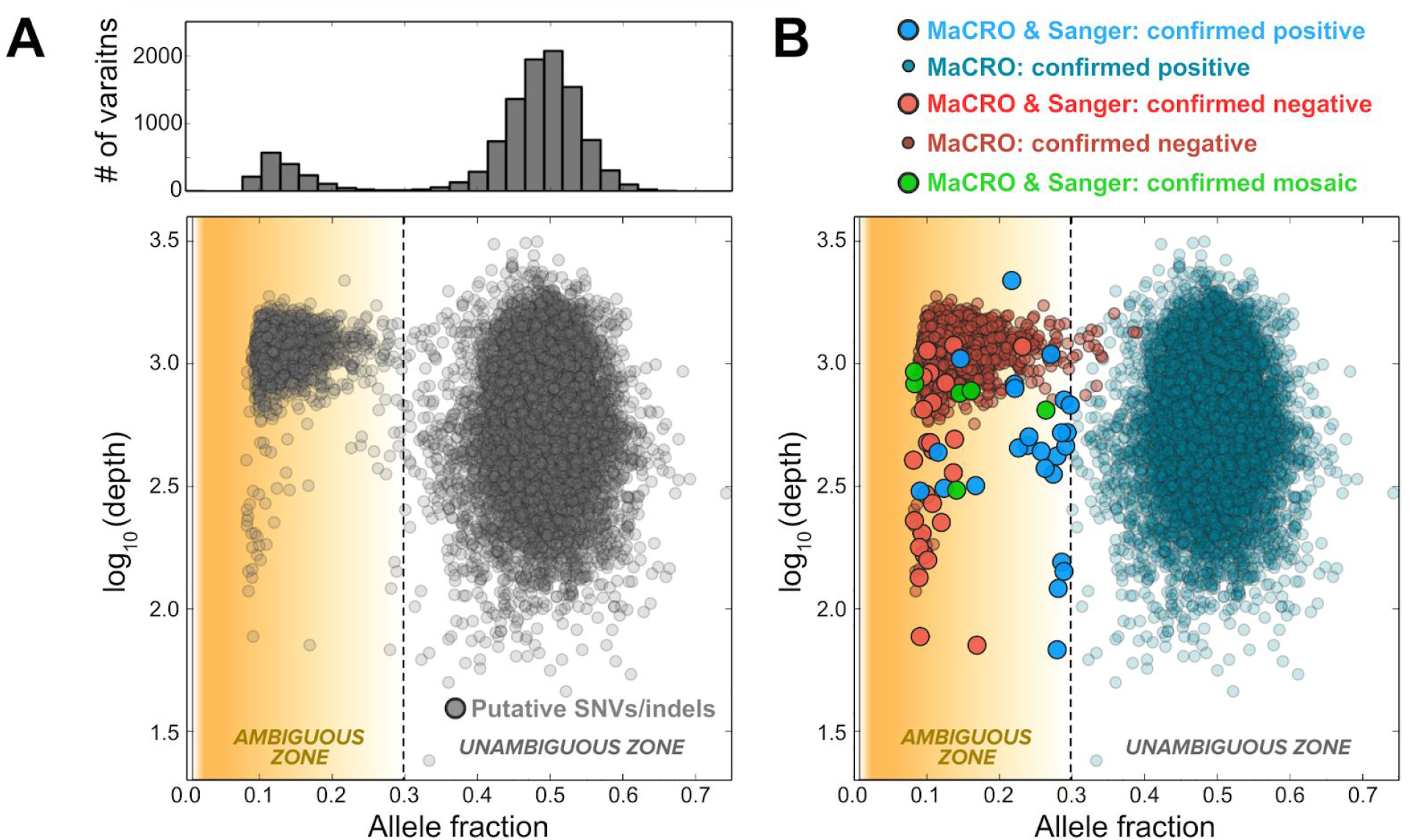
Ambiguous heterozygous variants with allele fraction can be resolved via MaCRO confirmation. **(A)** The bottom panel shows the distribution of putative positive calls (N = 11,099) as a function of allele fraction and the base-10 logarithm of NGS read depth fraction. A subset of calls populate an ambiguous zone where allele fraction is <30%. Each point in the scatter plot represents a single variant call. The top panel is a histogram of variant calls based on allele fraction alone. **(B)** MaCRO confirmation applied to the variant calls in (A) reveals a diversity of true genotypes in the ambiguous zone (see legend above plot). Sanger sequencing for samples selected from all unique, ambiguous variant sites upheld the MaCRO findings (large points in the scatter plot). Some MaCRO-confirmed negative calls have allele fraction >30%, illuminating how test accuracy could suffer if variant confirmation were constrained only to putative calls below a selected allele-fraction threshold.

### Software-assisted manual confirmation resolves ambiguous positive calls

MaCRO confirmation applied to NGS variant calls in the ambiguous zone revealed a mixture of positive, negative, and mosaic calls (Figure 2B). Though MaCRO was applied to all putative NGS positive calls in Figure 2B, we performed retrospective Sanger sequencing only on variants in the ambiguous zone (described below). We found that allele fraction and depth alone cannot resolve ambiguous variant calls, but software-assisted manual review of the NGS data was sufficient to yield a confident call without a requirement for Sanger confirmation.

Ambiguous calls were resolved by inspection of the raw NGS data during software-assisted manual confirmation. Figure 3 shows two variants from the study that are representative of others in the ambiguous zone: one is a confirmed heterozygous deletion with depressed allele fraction (Figure 3A-C), and the other is a site with spuriously elevated allele fraction at which the patient is confirmed to be homozygous for the reference allele (Figure 3D-F).

**Figure 3:**
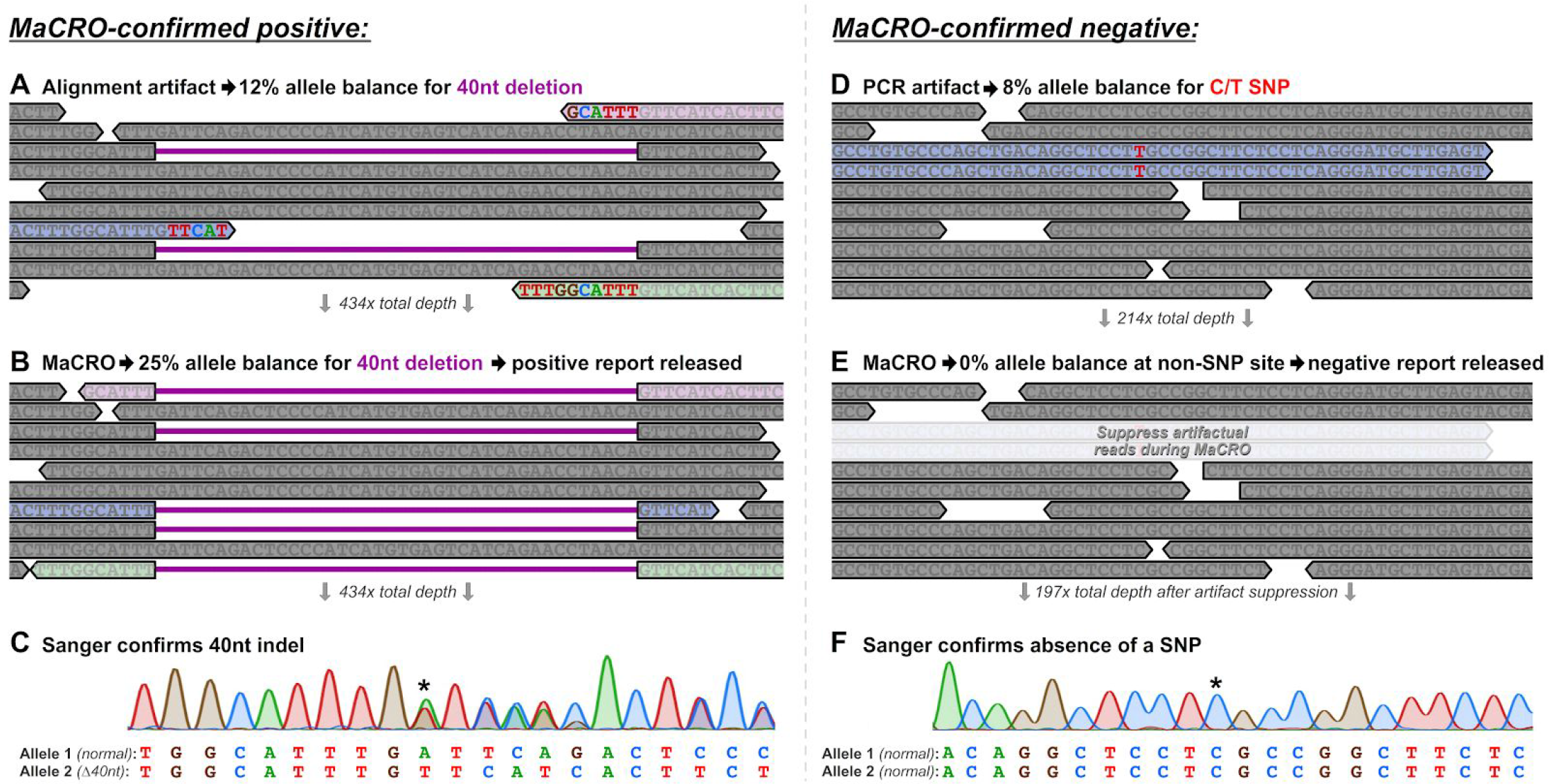
Examples of ambiguous positive variant calls resolved by MaCRO confirmation. **(A)** An excerpt of the NGS-read pileup in a sample with a 40nt deletion in *BRCA1* with 12% allele fraction reveals two reads spanning the deletion (reads with a purple gap). Of the 434x depth at this site, only 10x is depicted. The three pastel-shaded reads have mismatched bases at their 3’ ends, where the mismatched bases coincide with the deletion boundary, but the reads do not contribute to the allele fraction because the aligner does not recognize them as harboring the deletion. **(B)** MaCRO inspection of the pileup indicates that reads with mismatches are alignment artifacts and actually do have the 40nt deletion; properly viewing the reads as such effectively increases the allele fraction to 25% and raises the confidence of the heterozygous call. **(C)** Retrospective Sanger sequencing confirmed the existence of a heterozygous 40nt deletion; the asterisk indicates the deletion boundary. **(D)** A 10x excerpt from a 214x pileup of a putative SNV in *STK11* with 8% allele fraction shows that reads with the alternate base occur on the same strand and have the same endpoint. Alternate bases are not observed at any other location on either strand, suggestive of a PCR error (not shown is the fact that different probes captured fragments with the alternate base, consistent with the PCR error occuring during library preparation, not during the capture or sequencing process). **(E)** Suppressing the spurious reads during MaCRO evaluation reveals an otherwise clearly homozygous site. **(F)** At the site indicated with an asterisk, retrospective Sanger sequencing confirmed the homozygous-reference assertion from MaCRO.

The 40nt deletion in Figure 3A had low allele fraction due to an artifact of NGS-read alignment. Whereas the alignment software registered 12% of reads as harboring the deletion (i.e., those with purple lines in Figure 3A), the remaining reads were a mixture of reference-matching sequences and other reads that had a short series of SNVs near their termini (Figure 3A). Closer scrutiny of the pileup revealed that start and end points of the SNV series colocalized with the breakpoints of the deletion. Indeed, as shown in Figure 3B, the pastel-shaded soft-clipped reads were a perfect match with the 40nt deletion, but they were not initially aligned as such in the bioinformatics pipeline because they did not have enough sequence flanking the deletion. This expected limitation of the alignment software was overcome during MaCRO confirmation, enabling the reviewer to identify that the true allele fraction exceeded 12% and was indicative of the patient being heterozygous for a deleterious variant. Of the MaCRO-confirmed positive samples in the ambiguous zone (Figure 2B), 75% were indels (Table 1), and this example was the largest observed, which suggests efficacy of MaCRO across a range of indel sizes (Table 1).

**Table 1:**
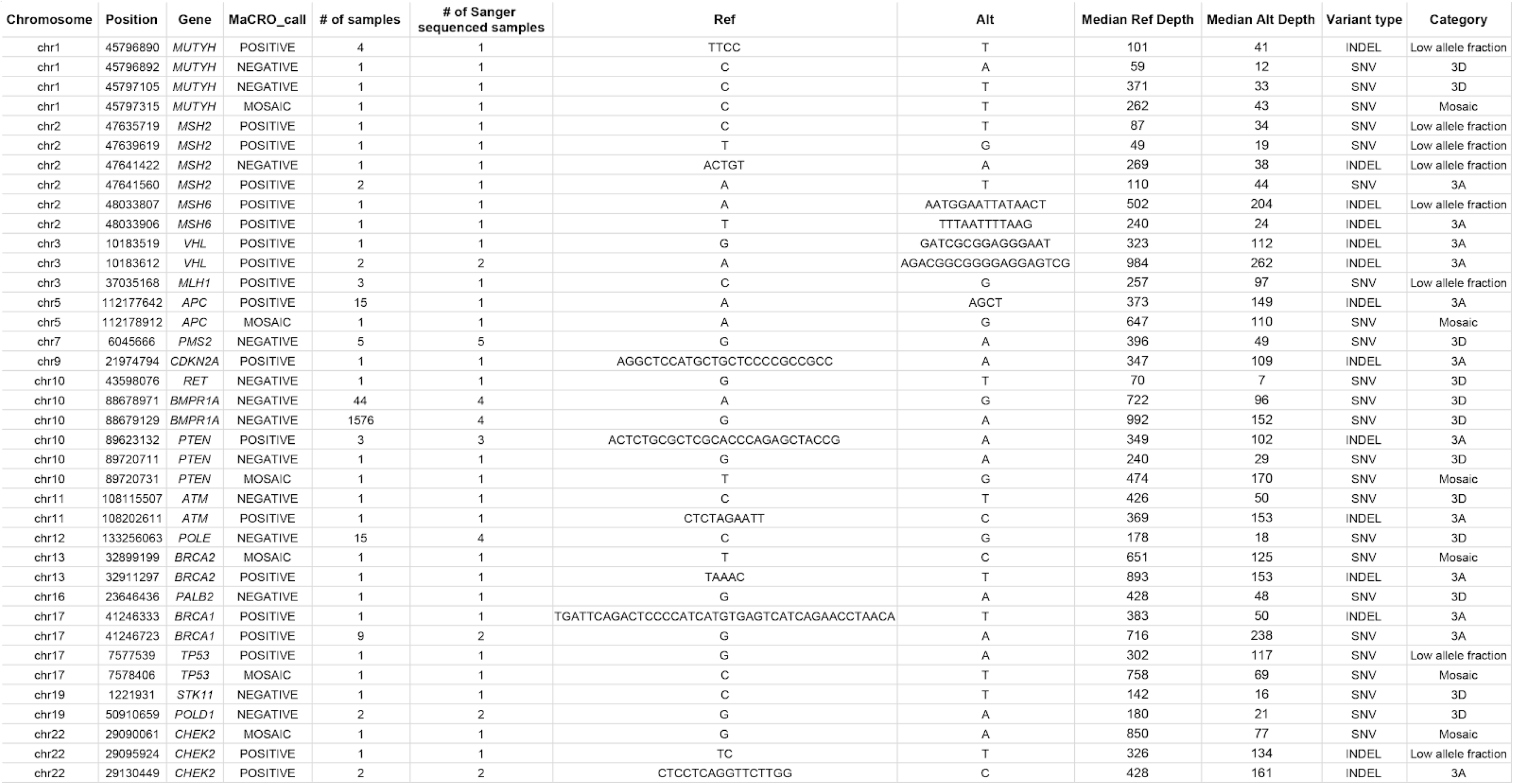
Overview of sites with ambiguous variant calls where samples underwent Sanger sequencing confirmation. The last column (“category”) indicates the four types of MaCRO data observed: “3A” and “3D” indicate NGS pileups with artifacts of the type shown in Figure 3A and Figure 3D, respectively; “Low allele fraction” indicates a pileup for a positive call where the allele fraction was very near 30%; some such sites are expected by chance given a large number of variant calls and intrinsic dispersion in the allele-fraction distribution (Figure 2). “Mosaic” indicates a pileup that resembles a typical heterozygous positive pileup but simply has depressed allele fraction.

Conspicuous PCR errors were another common artifact easily detectable during MaCRO-confirmation; one such error is illustrated in Figure 3D. In the pileup, the only evidence for heterozygosity was the T nucleotide, present in 8% of reads but always in reads that were from the same strand and at the same position. Using hybrid-capture technology, a real heterozygous site should have the alternate base interspersed among reads on both strands and with a range of endpoints (suggestive of being from different molecules in the genomic library). Further, because multiple capture probes interrogate each position and the capture probe for a particular fragment can often be inferred from paired-end data, MaCRO confirmation can also require that a legitimate variant be sampled via multiple probes. After suppressing the clearly spurious reads in the pileup (Figure 3E), the sample was MaCRO confirmed to be negative.

### Concordance between software-assisted manual confirmation and Sanger confirmation

We evaluated the efficacy of MaCRO confirmation by performing a retrospective Sanger sequencing analysis on variants in the ambiguous zone. All ambiguous variants (N=1,712) were concentrated at 39 sites (see Discussion). For all such sites, at least one sample with an ambiguous positive NGS call was Sanger sequenced (see Sanger sequencing results for each of the mosaic samples in Supplemental Figure S1). The genotypes elucidated via Sanger confirmation and MaCRO confirmation were perfectly concordant (shown comprehensively in Figure 4 and anecdotally in Figure 3C,F), indicating that Sanger sequencing offered no additional clinical benefit beyond the evaluation by MaCRO.

**Figure 4:**
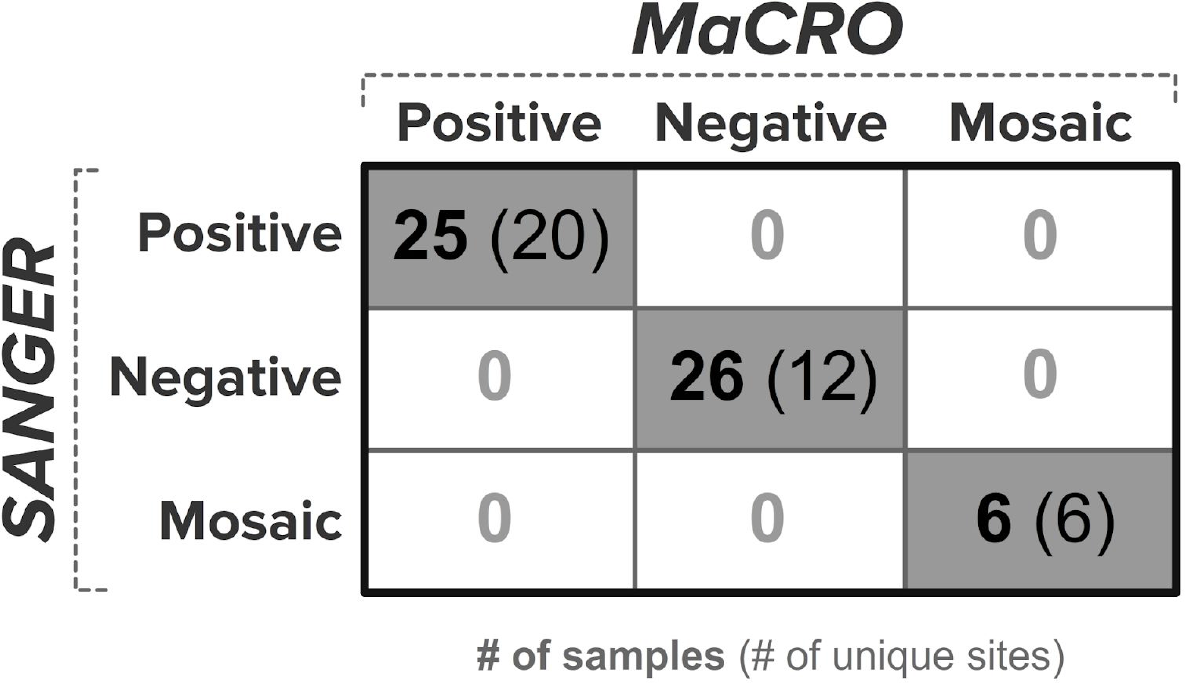
Sanger sequencing and MaCRO confirmation were concordant at all tested sites. In bold is the number of samples tested via both Sanger and MaCRO, and in parentheses is the number of unique sites.

### Feasibility of software-assisted manual confirmation in a high-throughput clinical laboratory

As genomic panels grow and screening becomes more widespread, it is important for confirmation methodologies to be efficient and scalable. We measured the time required for a single operator to apply MaCRO confirmation to a batch of samples (Figure 5A). Specifically, we queried the timestamp of every passed sample in 299 batches and calculated the duration between the first passed control and the third-to-last passed test sample, which was interpreted as the end point of MaCRO to account for rare samples that require retesting or dedicated follow-up attention from laboratory directors. The average MaCRO confirmation time is 15 minutes per batch (the mode is 8-12 minutes), and 96% of batches are reviewable by a single operator in less than one hour (Figure 5A). Therefore, with ~90 samples per batch, a single MaCRO reviewer can evaluate more than 1,000 samples with an expected 736 potentially reportable heterozygous calls in an eight-hour workday. Overall, 98.7% of HCS batches are MaCRO confirmed within 24 hours of completion of variant calling by the bioinformatics pipeline, reinforcing the rapidity of MaCRO confirmation (Figure 5B).

**Figure 5:**
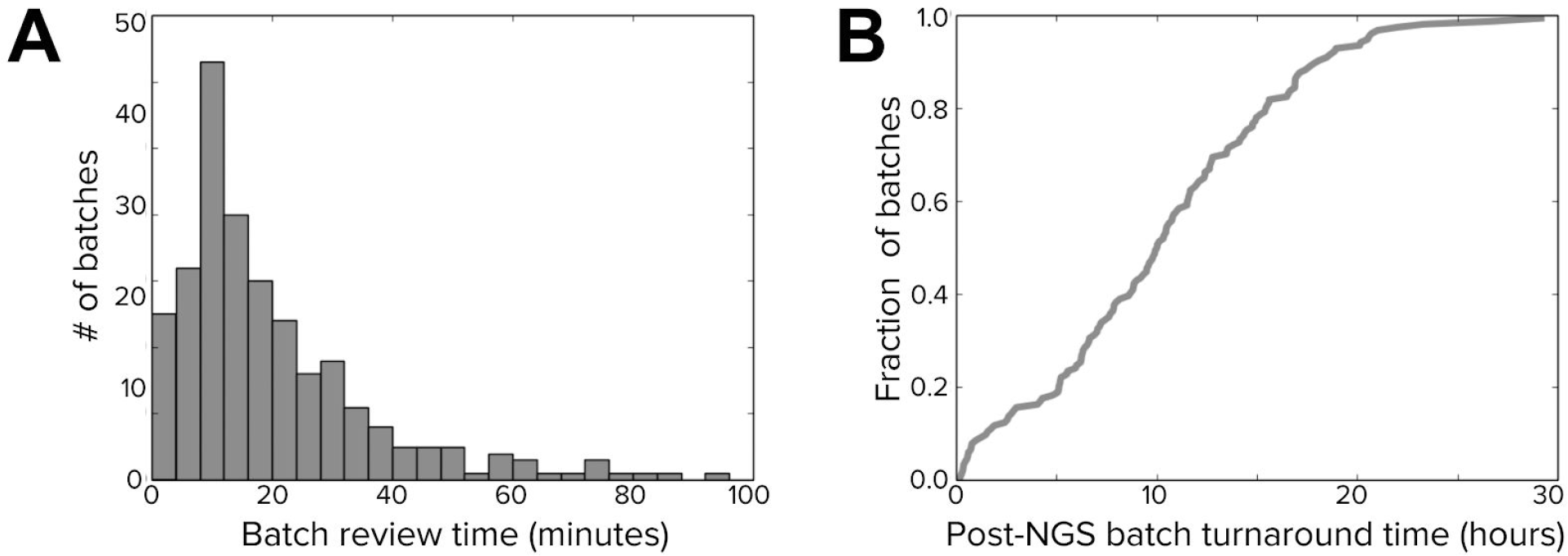
Time to perform MaCRO confirmation. **(A)** For 299 batches, the histogram shows the time between passing each batch’s first control sample and passing the third-to-last patient sample. **(B)** The cumulative-density plot shows the fraction of batches for which MaCRO confirmation occurred within the number of hours indicated (x-axis) following completion of NGS and bioinformatics processing.

## DISCUSSION

HCS results can inform major medical management decisions regarding cancer screening and prevention, so it is paramount for laboratories offering such testing to establish procedures that ensure high confidence in reported positive variants. Despite the high accuracy of automated variant calling from high-depth NGS data and modern bioinformatics pipelines, false positives are still possible, especially among low-confidence variant calls. Therefore, confirmation of positive calls emitted from an automated pipeline is clinically important, and Sanger sequencing is one such way for a laboratory to implement a confirmation workflow. However, it is not the only confirmation method, and here we have described and validated MaCRO confirmation as an effective alternative.

In a large patient cohort, MaCRO confirmation resolved genotypes among ambiguous low-allele-fraction calls, identifying them as positive, negative, or mosaic with the same accuracy as Sanger sequencing (notably, detection of mosaic variants is likely superior with MaCRO, as four of six mosaic calls detected by MaCRO had ≤15% allele fraction, near the limit of detection of Sanger sequencing alone^27^; Supplemental Figure S1). The 1,712 ambiguous calls we observed were concentrated among only 39 sites. This redundancy of many calls at a given site reinforces the idea that ambiguous calls are often systematic technical artifacts at the molecular level (e.g., elevated rate of PCR errors in homopolymers) or the algorithmic level (e.g., misalignment of large indels, alignment challenges in difficult-to-sequence regions like homopolymers, or spurious variants near the termini of reads). MaCRO was able to reveal these technical artifacts because the NGS chemistry enables parameterization of reference and alternate reads on three dimensions: the read strand, the fragment start position, and the likely capture probe for a fragment (inferred from paired-end data). Therefore, a stringent set of conditions could be applied to qualify a site as being heterozygous; it must have reference and alternate reads on both strands, at diverse positions, and from different capture probes. Conversely, spurious NGS calls could be identified by failing to show evidence of reference and alternate reads on any dimension (e.g., all alternate reads coming from one strand, one position, or one capture probe). Importantly, the high specificity conferred via MaCRO confirmation guards against false positives, which means that the bioinformatics pipeline can accordingly be optimized for sensitivity to minimize false negatives as well.

We observed that confirmation via software-assisted manual review and Sanger sequencing have comparable analytical performance in variant detection, but there are nontrivial differences in efficiency, scalability, affordability, and turnaround time. Each screen-positive sample receiving Sanger confirmation was estimated to incur one week of additional laboratory processing and a $240 cost^19^. By contrast, with an average MaCRO review time of 15 minutes for a batch of 90 samples, and assuming $100/hr compensation for a MaCRO operator, the average marginal review time and cost per sample are 10 seconds and $0.28, respectively. As a strategy to prevent ballooning the cost and turnaround time of HCS, several studies have suggested confining use of Sanger confirmation only to ambiguous calls and/or indels ^17,20,28^. But, MaCRO confirmation by comparison can plausibly be applied to all reportable calls in a clinical setting, which is important because we observed MaCRO-confirmed negatives with allele fraction >30% (Figure 2B). The increased speed and affordability of the MaCRO workflow relative to Sanger confirmation should increase accessibility of HCS testing and reduce overall turnaround time of the test, while simultaneously maintaining the quality of patient reports.

Although we performed Sanger confirmation on all ambiguous variant sites with allele fraction <30%, a limitation of our study is that we did not perform Sanger sequencing to verify MaCRO-confirmation performance for variant calls with >30% allele fraction. However, though we cannot disprove the possibility of a false positive among these confident calls, several studies have performed exhaustive Sanger sequencing on confident variants and found no false positives^17–19,21^. Another limitation is that, for individual sites at which multiple samples had the same variant (e.g., we found 15 variants in a *POLE* intron at position chr12:133256063), we often performed Sanger sequencing on one to five samples. Nevertheless, based on the perfect concordance observed between the two approaches at a variety of sites, we expect high performance of MaCRO in samples not further interrogated with Sanger sequencing. Finally, because MaCRO is human-operated, it is accordingly susceptible to human-operator error. We attempt to limit such error via engineering controls, e.g., by the interface requiring that any override to a call during MaCRO confirmation be certified by more than one MaCRO operator. However, a miscall could result if the initial reviewer did not appropriately interpret the data to conclude that a call should be overturned. Importantly, susceptibility to human error is not unique to MaCRO confirmation; it can equally impair interpretation of Sanger sequencing results.

We have demonstrated that software-assisted manual review of NGS data can be feasibly executed in a clinical setting and is highly accurate, yet two recent studies^17,18^ argue that Sanger confirmation is strictly required to resolve low-confidence NGS calls. To demonstrate their claim, both publications feature a figure highlighting the *MSH2* IVS5+3A>T SNV, a pathogenic variant at the boundary of a 27nt poly-A homopolymer. Despite the authors’ claim that Sanger sequencing is required to confirm this variant (correctly called in our HCS panel validation^6^ that utilized MaCRO), close visual inspection of the NGS pileups presented in their two publications—as would be done via MaCRO—clearly reveals the presence of the SNV. Therefore, had MaCRO confirmation been applied to the data presented, molecular confirmation via Sanger sequencing would be superfluous. Separately, neither study tabulates how many total variants confirmed via Sanger sequencing could not have been equivalently resolved from inspection of the NGS data itself. Without such an accounting, it is not clear that Sanger sequencing in particular is a necessity. Together, these publications underscore our assertion that confirmation of NGS-detected variants is strictly required, but the confirmatory method can take different forms.

MaCRO does not strictly supersede Sanger sequencing in our HCS testing. Sanger sequencing is used extensively during development and validation^6^ to characterize assay performance and identify potentially problematic regions. In production, however, the validated MaCRO protocol almost always provides sufficiently high confidence to issue a reported call. In cases where MaCRO was unable to resolve a particular variant, we pursued an alternative technique like Sanger sequencing. In sum, when carefully engineered, maintained, and validated, MaCRO confirmation, rather than Sanger confirmation, can be routinely used, with Sanger sequencing available as a secondary check only when needed.

Whether to use Sanger sequencing for confirmation of NGS-based results remains controversial, and medical societies have yet to issue a guideline regarding the use of Sanger sequencing. Our analysis of MaCRO confirmation demonstrates that there are multiple paths to achieve confident variant calls. Whereas Sanger sequencing has the undeniable virtue of being a classical and well-known technology, it remains only a proxy for quality testing, not a determinant. Because the sought-after goal is for genomic tests to be highly sensitive and specific, it should be the prerogative of the laboratory either to adopt a generally accepted practice like Sanger sequencing or to demonstrate openly that its alternative methodology yields comparably high-accuracy variant calls for patients. The work presented here shows that MaCRO confirmation does indeed yield comparably high-accuracy variant calls.

## ACKNOWLEDGEMENTS

We are grateful to Eerik Kaseniit, Katherine Johansen Taber, Kristin Price, Amy Sharma, Clement Chu, Imran Haque, and Eric Evans for support of the work and comments on the manuscript. Additionally, we thank the software engineering and CLIA teams at Counsyl for developing, operating, and supporting MaCRO for many years.

